# Projected spread of Zika virus in the Americas

**DOI:** 10.1101/066456

**Authors:** Qian Zhang, Kaiyuan Sun, Matteo Chinazzi, Ana Pastore-Piontti, Natalie E. Dean, Diana Patricia Rojas, Stefano Merler, Dina Mistry, Piero Poletti, Luca Rossi, Margaret Bray, M. Elizabeth Halloran, Ira M. Longini, Alessandro Vespignani

## Abstract

We use a data-driven global stochastic epidemic model to project past and future spread of the Zika virus (ZIKV) in the Americas. The model has high spatial and temporal resolution, and integrates real-world demographic, human mobility, socioeconomic, temperature, and vector density data. We estimate that the first introduction of ZIKV to Brazil likely occurred between August 2013 and April 2014 (90% credible interval). We provide simulated epidemic profiles of incident ZIKV infections for several countries in the Americas through February 2017. The ZIKV epidemic is characterized by slow growth and high spatial and seasonal heterogeneity, attributable to the dynamics of the mosquito vector and to the characteristics and mobility of the human populations. We project the expected timing and number of pregnancies infected with ZIKV during the first trimester, and provide estimates of microcephaly cases assuming different levels of risk as reported in empirical retrospective studies. Our approach represents an early modeling effort aimed at projecting the potential magnitude and timing of the ZIKV epidemic that might be refined as new and more accurate data from the region become available.

## Introduction

The Zika virus (ZIKV) is an RNA virus from first isolated in the Zika Forest of Uganda in the Flaviviridae family, genus Flavivirus [1, 2], 1947 [3]. It generally results in a mild disease characterized by low grade fever, rash, and/or conjunctivitis, though only approximately 20% of those infected are symptomatic [4]. Although there have been instances of sexual and perinatal/vertical transmission [5, 6, 7, 8, 9] and the potential for transmission by transfusion is present [10], ZIKV is spread primarily through infected *Aedes* mosquitoes [11, 12].

Until recently, ZIKV was considered a neglected tropical disease with only local outbreaks [13, 4, 14, 15]. In February 2015, the Brazilian Ministry of Health noted an unknown exanthematic disease spreading through six of its northeastern states [16]. The northeastern region of Brazil is characterized by high mosquito densities and dengue virus (DENV) transmission throughout the year. In mid May 2015, ZIKV was laboratory confirmed in multiple samples, establishing the presence of autochthonous transmission in Brazil. In September 2015, Brazil began to observe an increase in cases of microcephaly, a rare but serious neurologic birth defect resulting in a smaller than normal head [17]. On the basis of the reported microcephaly case cluster in Brazil and the previous association of ZIKV with Guillain-Barré syndrome (a post-infection autoimmune disease that affects the peripheral nervous system), the Director-General of the World Health Organization (WHO) declared a Public Health Emergency of International Concern (PHEIC) in February 2016 [18]. ZIKV has continued to spread throughout the Americas, with 47 countries and territories in the region reporting autochthonous transmission, excluding only Chile and Uruguay [19, 20]. Countries in the Americas reporting the highest incidence of ZIKV disease are also countries with historically endemic DENV transmission. As of recently, other countries with ZIKV outbreaks besides Brazil have reported cases of microcephaly and other birth defects associated with ZIKV infection during pregnancy (Zika Congenital Syndrome) [21].

While enhanced surveillance and new data have improved our understanding of ZIKV, many unknowns persist. There is uncertainty surrounding the time of introduction of the virus to the region, though early epidemiological and genetic findings estimate that ZIKV arrived in Brazil between May and December 2013 [22]. Furthermore, little is known about the spread of the virus in 2014 and 2015, prior to the WHO's alert in early 2016. Using a data-driven stochastic and spatial epidemic model, we analyze existing reports to gain insight into the first introduction into the region and early epidemic growth. We use the model to project future spatiotemporal spread in the Americas through February 2017, accounting for seasonal environmental factors and detailed population-level data. We also provide projections of the number of pregnancies infected with ZIKV during the first trimester along with estimates of the number of microcephaly cases per country using three different levels of risk based on empirical retrospective studies [23, 24].

## Results

### Introduction of ZIKV to the Americas

The precise location and start date of the outbreak are unknown. We identified twelve major transportation hubs in areas related to major events, such as the Soccer Confederations Cup in June 2013 and the Soccer World Cup in June 2014, both held in Brazil, and assumed a prior probability of introduction proportional to the daily passenger flow to each hub. We then considered introduction dates between April 2013 and June 2014, including the time frame suggested by phylogenetic and molecular clock analyses [22, 25] through the 2014 Soccer World Cup. Using Latin square sampling over this two-dimensional space, we calculated the likelihood of replicating the observed epidemic peak in Colombia (± 1 week), as reported by the Pan-American Health Organization (PAHO) [26], and the resulting posterior density of each location and date combination. The Colombian epidemic was used to calibrate this analysis because of the large number of cases observed and overall consistency in reporting.

In Fig. 1A we plot the posterior likelihood as a function of introduction date and location, and in Figs. 1B and 1C we plot the marginal posterior distributions of introduction date and location separately. The largest posterior density is associated with an introduction to Rio de Janeiro in December 2013. The 90% credible interval for the most likely date extends from August 2013 to April 2014, with its mode in December 2013. The most likely locations of ZIKV introduction, in descending order, are Rio de Janeiro (southeast), Brasilia (central), Fortaleza (northeast), and Salvador (northeast). While Rio de Janeiro experiences the greatest passenger flow, the city also experiences more seasonality in mosquito density making its likelihood to seed an epidemic sensitive to introduction date. The cities localized in the northeast of Brazil have less passenger flow compared to Rio de Janeiro but they have higher mosquito density and DENV transmission all year long. Brasilia, in comparison, has little seasonality in mosquito density and high traffic flow, though the area has low DENV transmission.

**Fig. 1:**
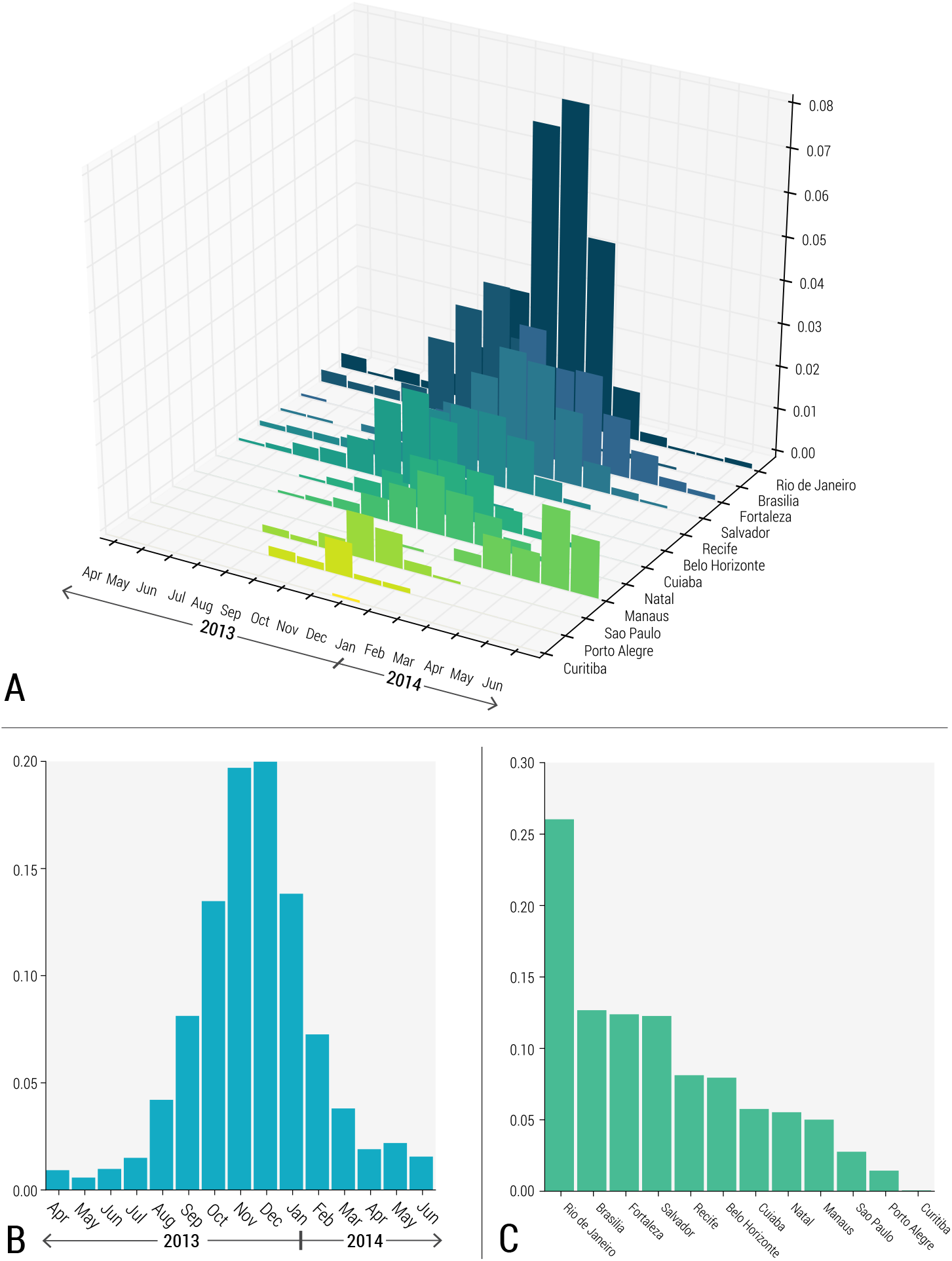
Posterior distribution for ZIKV introductions in twelve major transportation hubs in Brazil between April 2013 and June 2014, incorporating the likelihood of replicating the observed epidemic peak in Colombia. A) Full posterior distribution as a function of location and time of introduction. B) Marginal posterior distribution for time (month) of introduction. C) Marginal posterior distribution for location of introduction.

### Spatiotemporal ZIKV spread

Stochastic realizations satisfying the observed peak in Colombia define the model output used to project the spatiotemporal pattern of ZIKV spread in the Americas through February 2017. In Fig. 2 we plot the simulated epidemic profiles of incident ZIKV infections for several countries in the Americas, and in Tab. 1 we report the associated infection attack rates through February 1, 2016, when the WHO declared a PHEIC, and through February 28, 2017. The infection attack rate (infection AR) is defined as the ratio between the cumulative number of new infections during the considered period and the total population of a certain region. Estimates for additional countries in the Americas are available at the following webpage: www.zika-model.org. The earliest epidemic is observed in Brazil, followed by Haiti, Honduras, Venezuela and Colombia. The model projects that the epidemics in most of the countries would have declined around July 2016.

**Fig. 2:**
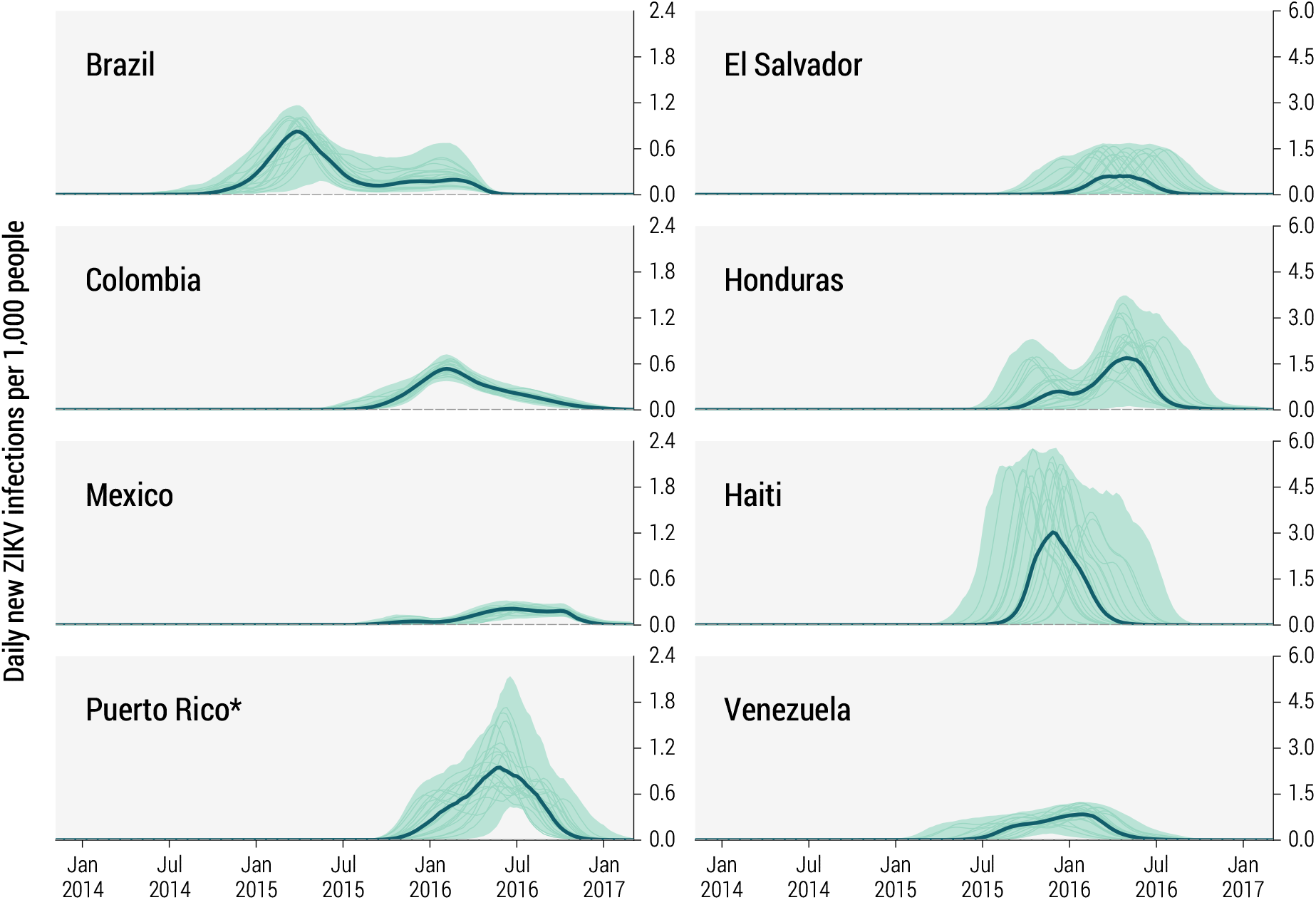
Estimated daily number of new ZIKV infections (per 1000 people) in eight affected countries in the Americas between January 2014 and February 2017. The bold line and shaded area refer to the estimated median number of infections and 95%CI of the model projections, respectively. Rates include asymptomatic infections. The median incidence is calculated each week from the stochastic ensemble output of the model and may not be representative of specific epidemic realizations. Thin lines represent a sample of specific realizations. *Puerto Rico curves are constrained under the condition that the peak of incidence curve is after March 1^*st*^ 2016, based on unpublished data [65].

**Tab 1.**
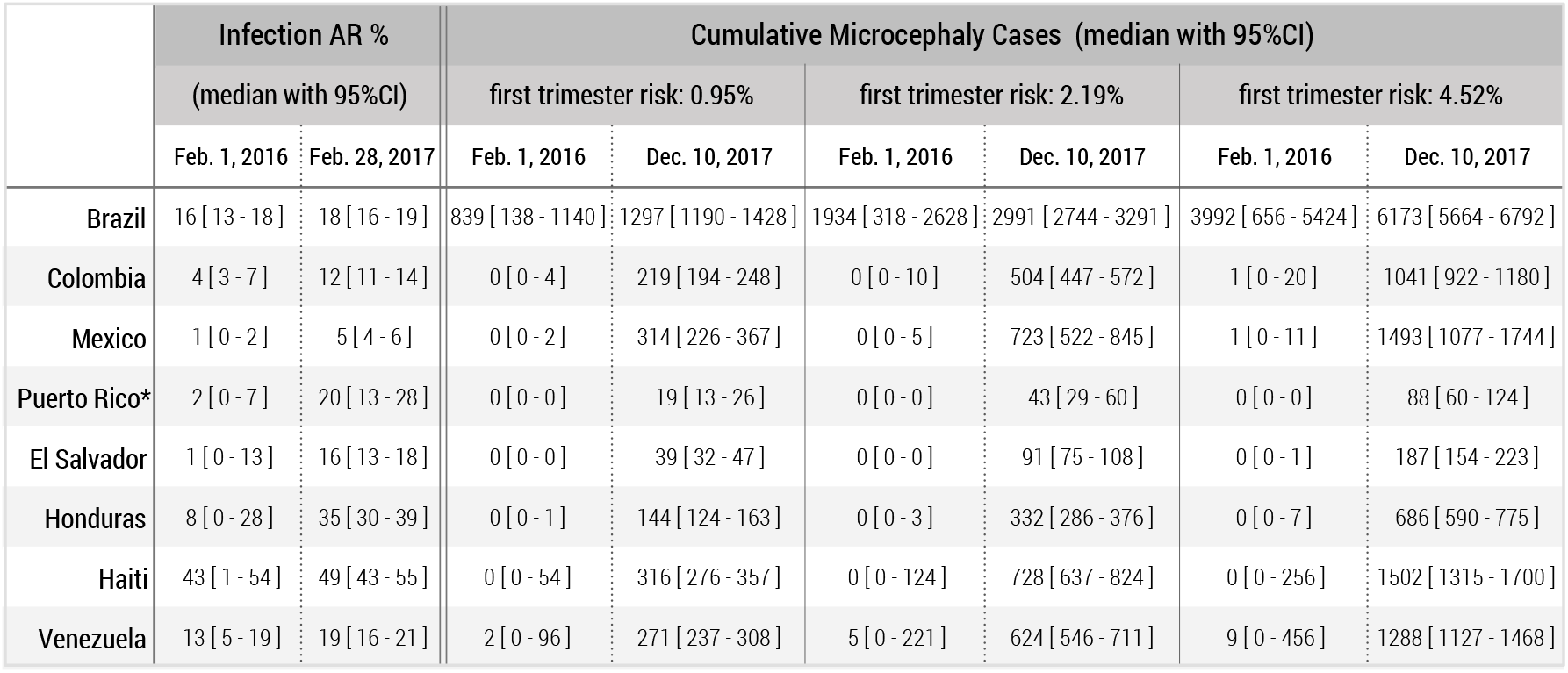
Projected ZIKV infection ARs through the time of the WHO declaration of a PHEIC on February 1, 2016, and through February 28, 2017, in eight affected countries in the Americas. Median estimates and 95%CIs are provided. ZIKV attack rates include asymptomatic infections. The denominator is the entire country population, including regions that are not exposed to the vector. Cumulative microcephaly cases due to ZIKV infection during the first trimester of pregnancy through the time of the WHO declaration of a PHEIC on February 1, 2016, and through December 10, 2017, in eight affected countries in the Americas. We consider three different risks of microcephaly associated with ZIKV infection during the first trimester: 0.95% first trimester risk based on a study of the 2013-2014 French Polynesian outbreak [23]; 2.19% (100% over-reporting) and 4.52% (no over-reporting) first trimester risks, based on a study of Bahia, Brazil [24], given a model-estimated 31% infection AR in Bahia. *Puerto Rico curves constrained under the condition that the peak of ZIKV incidence curve is after March 1^*st*^ 2016, based on unpublished data [65].

National infection ARs are projected to be high in Haiti, Honduras, and Puerto Rico. Countries with larger populations and more heterogeneity in mosquito density and vector borne disease transmission, such as Mexico and Colombia, experience much lower infection attack rates. For example, nearly half of Colombia's population resides in areas of high altitude where sustained ZIKV transmission is not possible. Because of the model's fine spatial and temporal resolution, we are able to observe significant variability in the ZIKV basic reproductive number *R*_0_ across locations and even within the same location at different times. These differences are driven by temperature, the vector distribution, and socioeconomic factors, among other variables. (Additional details are provided in the Materials and Methods.) In Fig. 3 we plot *R*_0_ in a number of areas at different times throughout the year. Equatorial regions experience less seasonality than non-equatorial regions where the changes in temperature have a strong impact on the mosquito population and, thus, *R*_0_. Large areas with unexposed populations are visible, such as in high altitude regions of Colombia.

**Fig. 3:**
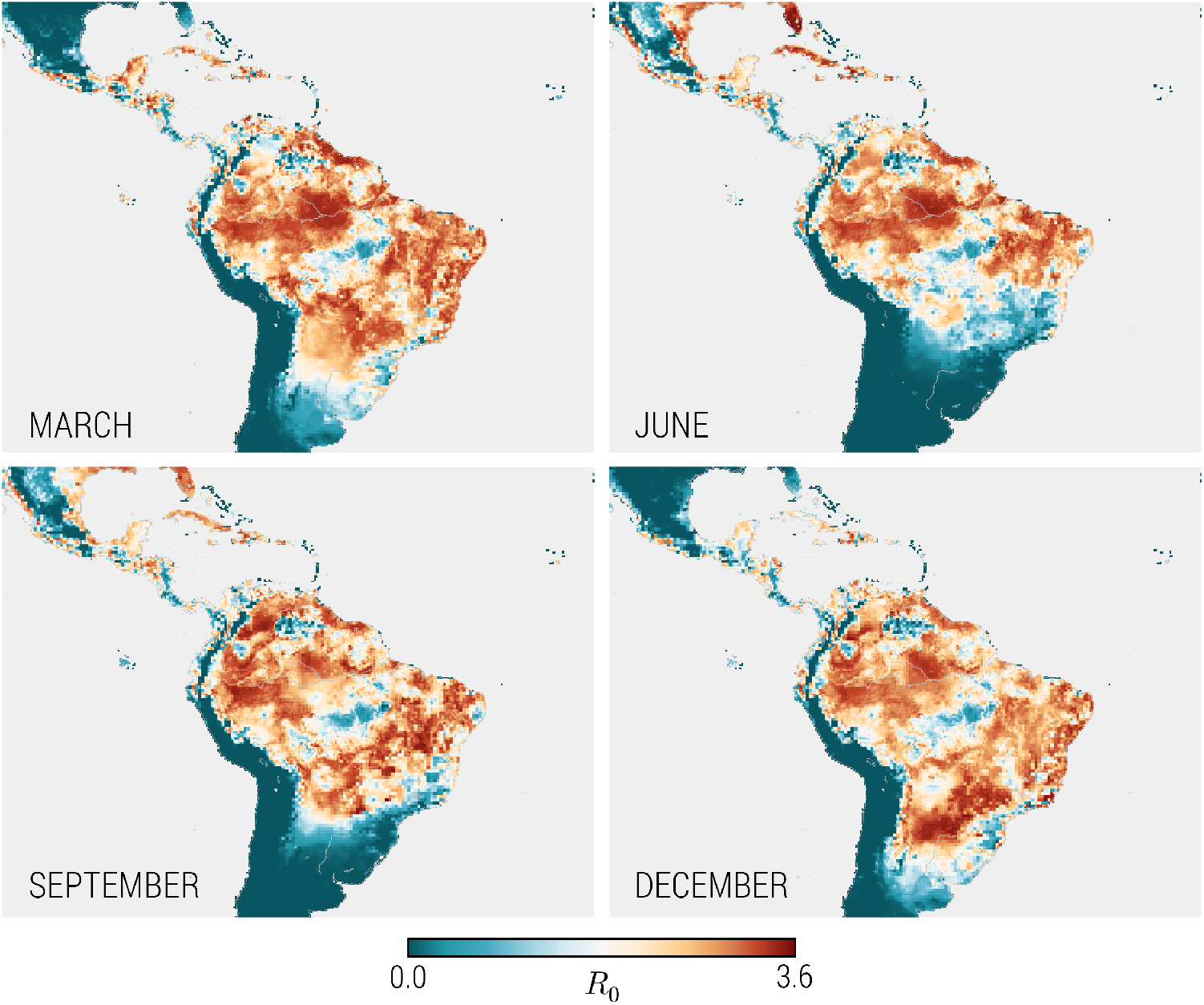
Monthly seasonality for the time- and location-dependent basic reproductive number, *R*_0_. The equatorial region displays less seasonality than the non-equatorial regions, where the changes of the season have a strong impact over the temperature and consequently over the basic reproductive number, *R*_0_.

### Projected ZIKV infections in childbearing women and microcephaly cases

Using the epidemic profiles generated by the model, we project the number of ZIKV infections in childbearing women following the model proposed in the study on ZIKV-microcephaly association of 2013-2014 French Polynesia outbreak [23]. In Fig. 4 we plot the daily number of births through December 2017 from women infected with ZIKV during the first trimester of pregnancy in several countries. Indeed, the first trimester of pregnancy is where the microcephaly risk is the highest [27, 23, 24]. The curves closely resemble the epidemic profiles in Fig. 2 but shifted forward in time by about 40 weeks. We construct our estimates using country-specific birth rates as detailed in Section 4 of the Supplementary Materials.

**Fig. 4:**
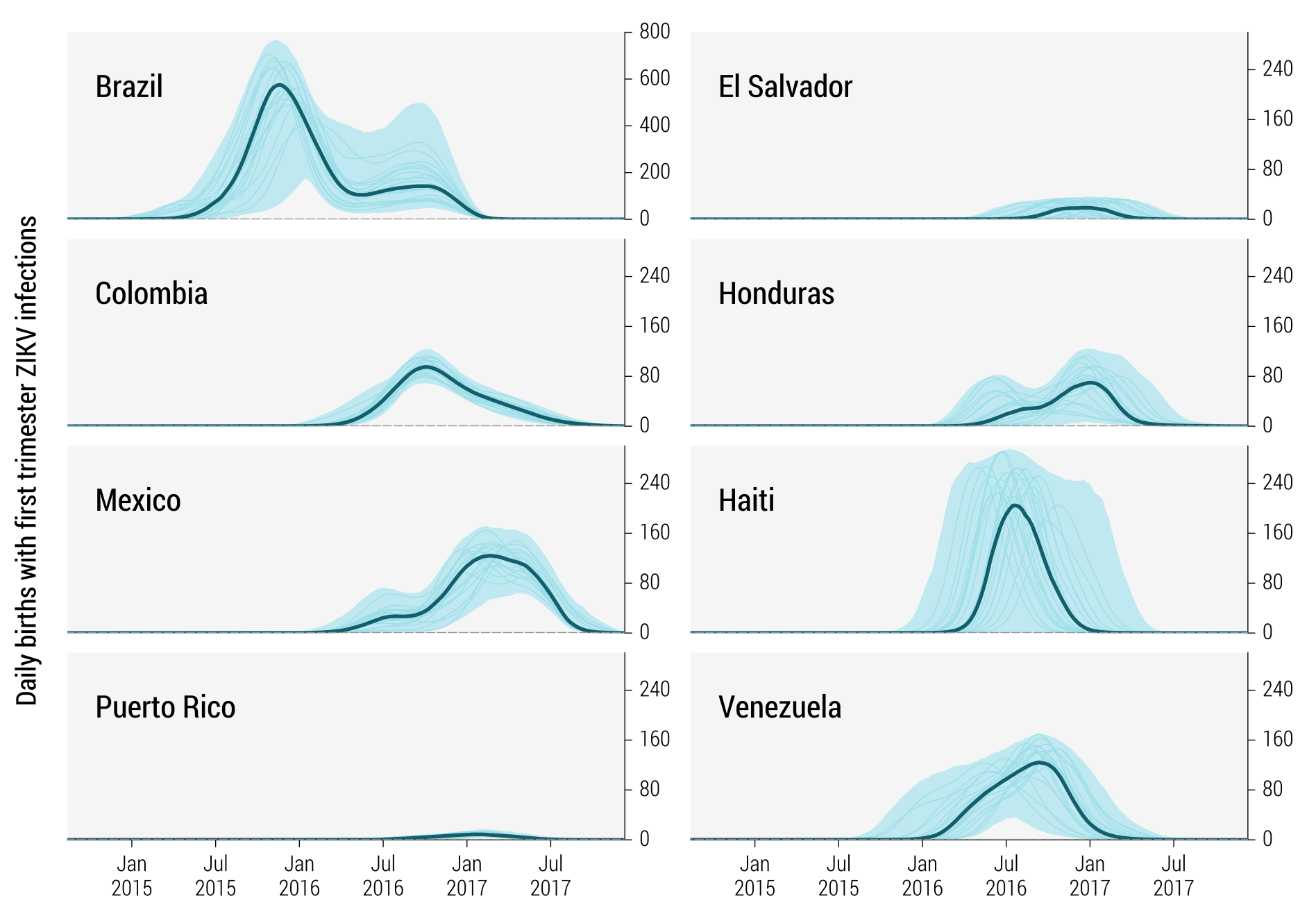
Estimated daily number of births between October 2014 and December 2017 from women infected with ZIKV during the first trimester of pregnancy in eight affected countries in the Americas. The bold line and shaded area refer to the estimated median number of births and 95%CI of the model projections, respectively. Note that Brazil is plotted with a different scale. The median curve is calculated each week from the stochastic ensemble output of the model and may not be representative of specific epidemic realizations. Thin lines represent a sample of specific realizations.

To estimate the number of microcephaly cases we adopt three different probabilities as reported in two empirical retrospectives studies [23, 24]. The first estimate of microcephaly risk for ZIKV infected pregnancies is 0.95% (95% confidence interval (CI) [0.34% – 1.91%]) from a study in French Polynesia [23]. The remaining two estimates come from a study performed in Bahia, Brazil [24]. Given an overall ZIKV infection AR of 31% (95% CI [26% – 36%]) in Bahia through February 2016, as determined by our model, the estimated first trimester microcephaly risks are 2.19% (95% CI [1.98% – 2.41%]), assuming 100% over-reporting of microcephaly cases, and 4.52% (95% CI [4.10% – 4.96%]), assuming no over-reporting. The estimates do not account for miscarriages or other complications occurring during pregnancy.

In Tab. 1 we report the projected cumulative number of microcephaly cases up to February 1, 2016, and to December 10, 2017. By the time the WHO declared a PHEIC, Brazil was the only country with a substantial (>100) number of ZIKV-attributable microcephaly cases, with cases expected to appear through July 2017. For Colombia, the model projects the largest number of new microcephaly cases between October and December 2016. In Venezuela, the peak in microcephaly cases is projected to have occured in September/October 2016, continuing through February 2017. In Puerto Rico the largest number of microcephaly cases is expected to occur in January 2017.

### Sensitivity to mosquito vector

Simulations reported here consider both *Ae. aegypti* and *Ae. albopictus* as competent ZIKV vectors, though less is known about the vectorial capacity of *Ae. albopictus*. A sensitivity analysis considering *Ae. aegypti* as the only competent vector is provided in the Supplementary Materials with all figures and tables replicated for this scenario. Overall, results are similar as transmission due to *Ae. aegypti* increases to compensate for the absence of the other vector. Differences in the infection ARs, however, are observed in areas where *Ae. albopictus* is the most common or the only vector. For example, the infection AR in Brazil through February 28, 2017, decreases from 18% (95% CI [16% – 19%]) to 16% (95% CI [14% – 17%]) if we consider only *Ae. aegypti*. During the same time period, the infection AR in Mexico decreases from 5% (95% CI [4% – 6%]) to 4% (95% CI [2% – 5%]).

### Model validation

Several approaches were used to independently validate the model with data not used for calibration, as shown in Fig. 5. In Fig. 5A, we compare model-based projections of the number of ZIKV infections by states in Colombia with observed surveillance data through October 1, 2016 [26]. As expected for a typically asymptomatic or mild disease, the model projects a much larger number of infections than that captured by surveillance, suggesting a reporting and detection rate of 1.02% ± 0.93% (from linear regression analysis). However, the observed data and model estimates are well-correlated (Pearson's *r* = 0.68, *p* < 0.0001), replicating the often several orders of magnitude differences in infection burden across states within the same country.

**Fig. 5:**
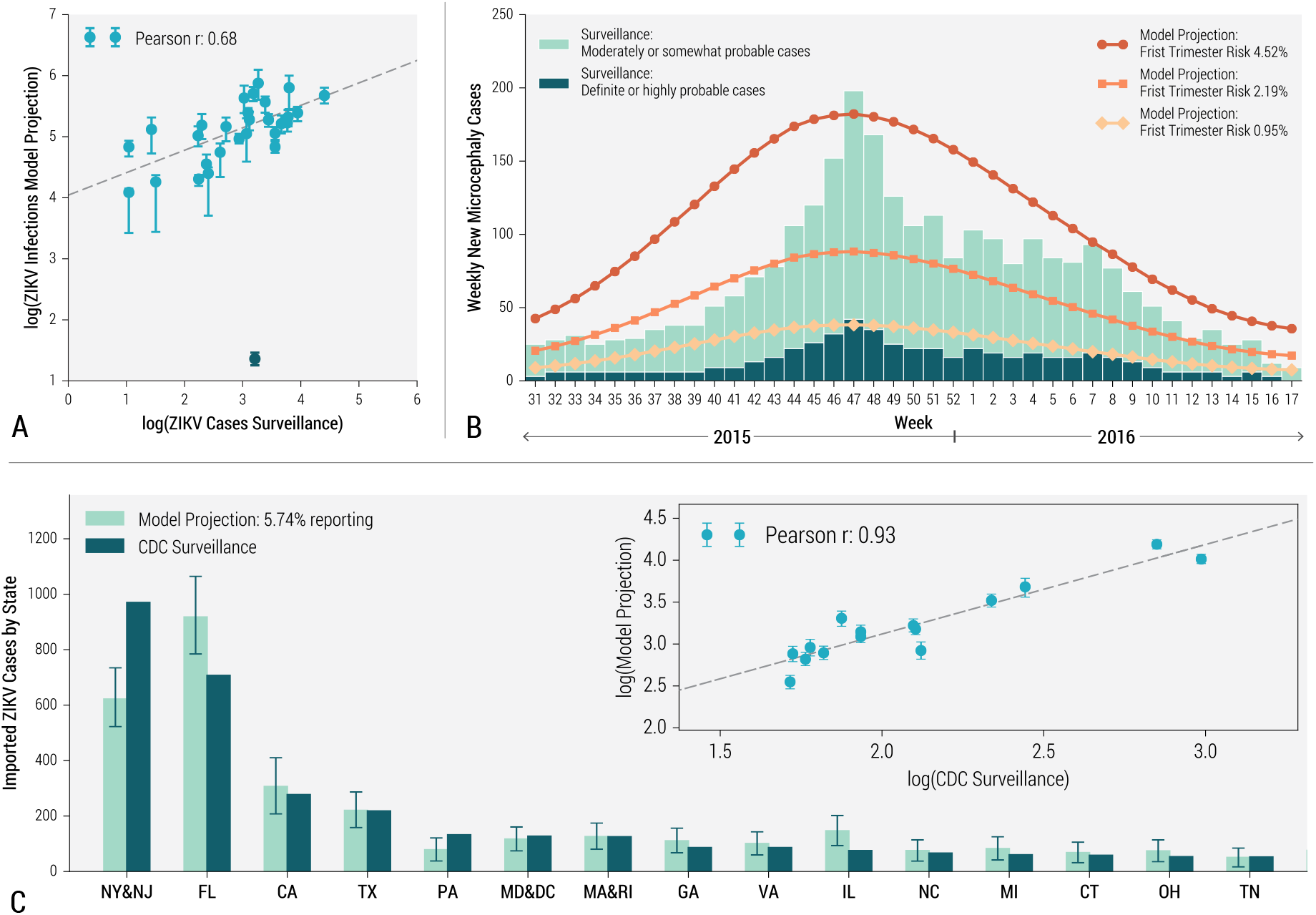
A) Correlation between the number of ZIKV cases by state in Colombia as reported by surveillance data through October 1, 2016 [26], compared with state-level model projections of infections (median with 95% CI). Pearson's *r* correlation coefficient is reported for the linear association on the log scale. The outlier (in dark green) excluded from the statistical analysis corresponds to the Arauca region. B) Timeline of microcephaly cases in Brazil through April 30, 2016. Bar plot shows weekly definite (or highly probable cases) and moderately (or somewhat probable cases) from surveillance data [28]. Line plots indicate estimated weekly new microcephaly cases given three levels of first trimester risk: 4.52% (round) [24], 2.19% (square) [24], and 0.95% (diamond) [23]. C) Bar plot of ZIKV infections imported into the continental USA by state(s) as reported by CDC surveillance through October 5, 2016 [29], and compared to model projections (median with 95% CI) for the same period assuming 5.74% reporting/detection. The insert shows the correlation between CDC surveillance data and model projections (median with 95%CI).

In Fig. 5B we compare observed data on weekly counts of microcephaly cases reported in Brazil through April 30, 2016 [28] with estimates from the model for each projected level of microcephaly risk given first trimester ZIKV infection. The three model projection curves vary in magnitude but replicate peaks consistent with the observed data. As the fraction of cases that are confirmed in Brazil is relatively low, it is not possible to identify the most likely level of risk, though the plot suggests that the risk might exceed the lowest estimate of 0.95% [23].

As the computational approach explicitly simulates the number of daily airline passengers traveling globally, the microsimulations allow us to track ZIKV infections imported into countries with no autochthonous transmission. In Fig. 5C we plot the number of importations into states in the continental United States (USA) through October 5, 2016, as reported by the Centers for Disease Control and Prevention (CDC) [29] and compare these results with model projections. As the detection rate of ZIKV infections is very low, there are significantly fewer reported cases than projected; we estimate through a linear regression fit that 5.74% ± 1.46% of imported infections are detected. Nonetheless, the model projections are highly correlated with the observed data (Pearson's *r* = 0.93, *p* < 0.0001), as shown in the inset of Fig. 5C. A further validation of the model is provided by the reported number of ZIKV cases of pregnant women in the USA. A high detection rate is expected in this closely monitored population. As of September 29, 2016, 837 pregnant women in the continental USA were laboratory-confirmed for ZIKV, all of whom were imported cases. Since pregnant women comprise approximately 1% of incoming airline traffic flow from the rest of the Americas [30], one can roughly estimate 83, 700 infections, which is in the ballpark of our modeling results projecting 57, 910 (95% CI [50,138 – 66,608]) infections imported into the USA by early October 2016. These results are relevant for ZIKV risk assessment in USA [31, 32].

## Discussion

We use computational modeling to reconstruct the past and project the future spatiotemporal spread of ZIKV in the Americas. To identify the likely date and location of ZIKV's first introduction to the Americas, posterior densities were estimated for twelve major travel hubs in Brazil over a range of dates. The marginal posterior distributions suggest an introduction between August 2013 and April 2014 in a number of potential locations, including Rio de Janeiro, Brasilia, Fortaleza, Salvador. This date range partially overlaps with that suggested by a recent phylogenetic analysis [22, 25], though ours includes later potential introductions.

We provide epidemic curves for incident ZIKV cases in eight affected countries in the Americas. The results obtained are in good agreement with model-based projections achieved with a different approach as described elsewhere [33]. Though the initial introduction of ZIKV could date back to August 2013, most countries do not experience the first wave of the epidemic until the early months of 2016. Brazil is the only country that appears to have a well-defined first peak in March 2015, consistent with reports from the northeast region [34]. The model suggests two epidemic waves in Brazil. The first wave, occurring between January and July 2015, corresponds to early outbreaks in the northeast region (Maranhao, Bahia, and Rio Grande do Norte) and later on to the rest of the country. This first wave was not recognized as ZIKV until early 2016. The second wave, between January and May 2016, affects mostly southern states in Brazil [35].

The virus also circulated early on in the Caribbean, with ZIKV samples isolated in Haiti at the end of 2014 and a possible first peak occurred in October 2015 [36]. Colombia first isolated ZIKV in October 2015, at which time it spread rapidly from the Caribbean coast to cities infested with *Ae. aegypti* [37]. The model suggests an introduction to Colombia as early as March-April 2015, potentially overlapping with the Easter holiday, which is a period of high mobility within and between countries. ZIKV transmission in Venezuela follows a similar trajectory, first isolated in November 2015 and present in all states by July 2016 [38]. Since March 2016, reported cases have declined in both countries, consistent with our model estimates.

Our model estimates ZIKV transmission in El Salvador and Honduras increasing around July 2015. ZIKV was first detected in El Salvador in November 2015 and in Honduras in December 2015 [39, 40]. Though the first ZIKV infection was confirmed in Puerto Rico in the last week of December 2015 [41], the model estimates ZIKV transmission in Puerto Rico beginning around August 2015. In Mexico, the first infection was reported to the surveillance system at the end of November 2015 [42], though circulation may have begun in September 2015.

Another prominent feature emerging from the numerical results is the extreme heterogeneity in the infection ARs across countries. We find more than a sevenfold difference between Honduras and Mexico, exhibiting infection ARs of 35% (95% CI [30% – 39%]) and 5% (95% CI [4% – 6%]), respectively. These large differences in infection ARs, which are observable at finer geographical resolutions as well, stem from variation in climatic factors, mosquito densities, and socioeconomic variables.

The epidemic has moved slowly and is mostly constrained by seasonality in ZIKV transmissibility. Seasonal drivers and time of introduction result in multiple waves [43] across several countries, as projected for Brazil, Honduras, and Mexico. Notably, though, incidence rates dramatically decrease by the end of 2016 in all countries described. These results suggest that, after the initial wave in an almost fully susceptible population, ZIKV epidemics could settle into the typical seasonal pattern of mosquito borne diseases like DENV. Transmission may be low for several years with a gradual build-up in susceptibility due to births [44]. Precise projection of long-term ZIKV transmission is very important to plan for future Zika control activities and for finding sites for phase III Zika vaccine trials. This is a topic for future research.

We project the numbers of births from women who were infected with ZIKV during the first trimester of their pregnancy. There is a well-defined time lag between the epidemic curve and this birth curve. Brazil, that likely experienced its first ZIKV epidemic peak in March 2015, had a sharp rise in microcephaly cases in September 2015, consistent with what was observed in the field [28]. Venezuela also experienced a comparatively early peak, but it is possible that this has not been reported because case detection and reporting are low due to the political climate. Our estimates correlate with the situation in Colombia where 57 confirmed cases of congenital Zika syndrome (373 additional cases are under study) have been observed as of October 29, 2016 [45]. Note that the projected number of microcephaly cases estimated by the model varies considerably depending on the assumed first trimester risk, for which only retrospective estimates are available [23, 24]. We also note that with as high as 80% of ZIKV infections being asymptomatic [4, 27], most of the ZIKV-infected pregnant women giving birth may not have experienced symptoms during pregnancy. Thus, clinicians should be cautious before ruling out ZIKV as the cause of birth defects. The results presented here, however, could be used as a baseline to uncover possible disagreement with the observed data and highlight the need of additional key evidence for our understanding of the link between ZIKV and neurologic birth defects [46].

Available data on the ZIKV epidemic suffer from several limitations. Although the disease has likely been spreading in the Americas since late 2013, infection detection and reporting began much later and likely increased after the WHO's declaration of a PHEIC in February 2016. Case reporting is inconsistent across countries. Furthermore, comparatively few infections are laboratory-confirmed; this presents an additional challenge as symptomatic cases with other etiologies may be misdiagnosed, and asymptomatic infections are almost entirely missed. Once a reliable ZIKV antibody test is available, seroprevalence studies can help determine the full extent of these outbreaks. For external validation, we compare modeling results with data from Brazil, Colombia, and the USA that were not used to calibrate the model. We are able to replicate relative trends, though we estimate significantly higher absolute numbers, suggesting reporting and detection rates of about 6%.

The modeling approach presented here is motivated by the need for a rapid assessment of the ZIKV epidemic, and it contains assumptions and approximations unavoidable due to the sparsity of available data. As a result, transmission is modeled assuming ZIKV behaves similarly to DENV and other mosquito-borne diseases, and further research is needed to provide ZIKV-specific parameter estimates. Mosquito presence maps were available from published data but have limitations as detailed in the literature. Sexual and other modes of transmission were not incorporated in the model. The specific socio-economic features of airline travelers was also not included. Finally, we do not model public health interventions to control the vector population or behavioral changes due to increased awareness. These results may change as more information becomes available from ZIKV affected regions to refine the calibration of the model.

## Conclusions

The model presented here provides an early methodological framework for the analysis of the global spread of ZIKV. The model captures the slow dynamic of the epidemic characterized by heterogeneity in the infection AR as well as the temporal pattern resulting from local weather, population-level characteristics, and human mobility. Although the modeling results should be interpreted cautiously in light of the assumptions and limitations inherent to the approach, the framework emerging from the numerical results may help in the interpretation of data and provide indications of the magnitude and timing of the epidemic, as well as aid in planning for international and local outbreak response, and for the planning of phase III vaccine trial sites.

## Materials and Methods

### Model summary

To study spatiotemporal ZIKV spread, we use the Global Epidemic and Mobility Model (GLEAM), a previously described individual-based, stochastic and spatial epidemic model [47, 48, 49, 50, 51, 52]. This model integrates high-resolution demographic, human mobility, socioeconomic [53], and temperature data [54]. It was expanded to incorporate data on *Aedes* mosquito density [55] and the association between socioeconomic factors and population risk of exposure [33, 56]. Similar to previous arbovirus modeling approaches [15], we use a compartmental classification of the disease stages in the human and mosquito populations, assigning plausible parameter ranges based on the available ZIKV literature and assumed similarities between ZIKV and DENV.

### Global model for the spread of vector-borne diseases

The GLEAM model is a fully stochastic epidemic modeling platform that utilizes real world data in order to perform in-silico simulations of the spatial spread of infectious diseases at the global level. GLEAM uses population information obtained from the high-resolution population database of the *Gridded Population of the World* project from the Socioeconomic Data and Application Center at Columbia University (SEDAC) [57]. The model considers geographical cells of 0.25 × 0.25 degree, corresponding to an approximately 25 × 25 km square for cells along earth's Equator. GLEAM groups cells into subpopulations defined by a Voronoi-like tessellation of the earth's surface centered around major transportation hubs in different urban areas. The model includes over 3,200 subpopulations in about 230 different countries (numbers vary by year).

Within each subpopulation, a compartmental model is used to simulate the disease of interest. The model uses an individual dynamic where discrete, stochastic transitions are mathematically defined by chain binomial and multinomial processes. Subpopulations interact through the mechanistically simulated mobility and commuting patterns of disease carriers. Mobility includes global air travel [58], and GLEAM simulates the number of passengers traveling daily worldwide using available data on origin-destination flows among indexed subpopulations.

The transmissibility of vector-borne diseases is associated with strong spatial heterogeneity, driven by variability in vector abundance and characteristics of the exposed populations. Many locations, such as those at high elevation, are not at risk of autochthonous ZIKV transmission simply because the vector is absent. In other locations, the vector may be present but sustained transmission is not possible because of environmental factors that affect the vector's population dynamics, such as temperature or precipitation. Housing conditions, availability of air-conditioning, and socioeconomic factors also contribute significantly to determining the fraction of the population likely exposed to the vector. To extend the GLEAM model to simulate vector-borne diseases, a number of new data sets with high spatial resolution were integrated, including the following:

- *Global terrestrial air temperature data*: The global air temperature data set [54] contains monthly mean temperatures at a spatial resolution of 0.5 × 0.5 degree. To match the spatial resolution of GLEAM's gridded population density map, the temperature for each population cell is extracted from the nearest available point in the temperature data set. Daily average temperatures were linearly interpolated from each population's monthly averages.
- *Global Ae. aegypti and Ae. albopictus distribution*: The Global *Ae. aegypti* and *Ae. albopictus* distribution database provides uncertainty estimates for the vector's distribution at a spatial resolution of 5 × 5 km [55].
- *Geolocalized economic data*: The geophysically scaled economic dataset (G-Econ), developed by Nordhaus et.al. [59], maps the per capita Gross Domestic Product (computed at Purchasing Power Parity exchange rates) at a 1 × 1 degree resolution. To estimate the per capita Gross Cell Product (GCP) at Purchasing Power Parity (PPP) rates, the amount is distributed across GLEAM cells proportionally to each cell's population size. The data have also been rescaled to reflect 2015 GDP per capita (PPP) estimates.

These databases are combined to model the key drivers of ZIKV transmission, as illustrated in combination with necessary parameters in Fig. 6. Temperature impacts many important disease parameters, including the time- and cell-specific values of *R*_0_ whose variation induces seasonality and spatial heterogeneity in the model. Temperature data are also used together with the mosquito presence distribution data to define the daily mosquito abundance (number of mosquitoes per human) in each cell as detailed in Sec. 2 of the Supplementary Materials. Data on mosquito abundance and temperature are used to identify cells where ZIKV outbreaks are not possible because of environmental factors. The human population in these cells is thus considered unexposed to ZIKV and susceptible individuals are assigned an environmental rescaling factor, *r_env_*, as described in Sec. 3 of the Supplementary Materials. Finally, we use historical data and G-Econ to provide a socioeconomic rescaling factor, *r_se_*, reflecting how exposure to the vector is impacted by socioeconomic variables such as availability of air-conditioning; the derivation of this rescaling factor is provided in Sec. 3 of the Supplementary Materials.

**Fig. 6:**
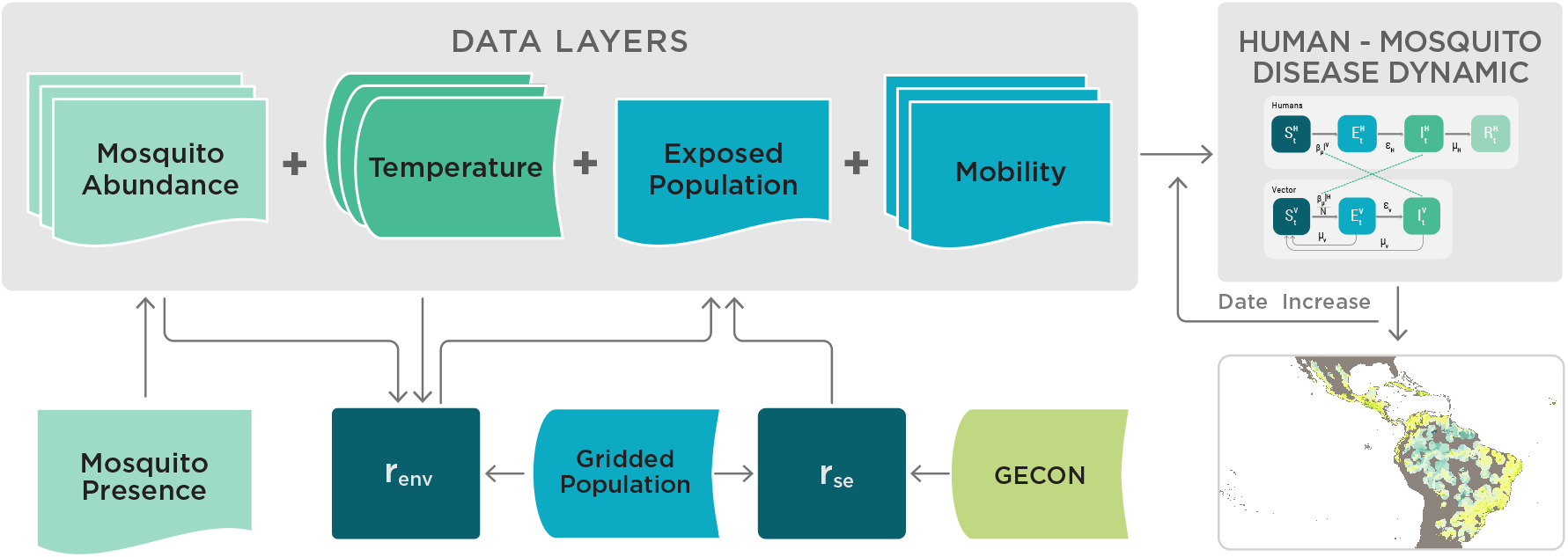
Schematic representation of the integration of data layers and the computational flow chart defining the Global Epidemic and Mobility (GLEAM) model for ZIKV.

Once the data layers and parameters have been defined, the model runs using discrete time steps of one full day to run the transmission dynamic model (described in detail below), incorporate human mobility between subpopulations, and partially aggregate the results at the desired level of geographic resolution. The model is fully stochastic and from any nominally identical initialization (initial conditions and disease model) generates an ensemble of possible epidemics, as described by newly generated infections, time of arrival of the infection in each subpopulation, and the number of traveling carriers. The Latin square sampling of the initial introduction of ZIKV in Latin America and the ensuing statistical analysis is performed on 150,000 stochastic epidemic realizations. The sensitivity analysis considers a total of 200, 000 more realizations.

### ZIKV transmission dynamics

Fig. 7A describes the compartmental classifications used to simulate ZIKV transmission dynamics. Humans can occupy one of four compartments: susceptible individuals *S^H^* who lack immunity against the infection, exposed individuals *E^H^* who have acquired the infection but are not yet infectious, infected individuals *I^H^* who can transmit the infection (may or may not display symptoms), and removed individuals *R^H^* who no longer have the infection. We consider the human population size to be constant, *i.e*. *S^H^*+*E^H^*+*I^H^* + *R^H^* = *N^H^*. The mosquito vector population is described by the number of susceptible *S^V^*, exposed *E^V^*, and infectious mosquitoes *I^V^*. The transmission model is fully stochastic. The transitions across compartments, the human-to-mosquito force of infection, and the mosquito-to-human force of infection are described by parameters that take into account the specific abundance of mosquitoes and temperature dependence at the cell level. Exposed individuals become infectious at a rate *ε_H_* which is inversely proportional to the mean intrinsic latent period of the infection [60]. These infectious individuals then recover from the disease at a rate *μ_H_* [15] which is inversely proportional to the mean infectious period. The mosquito-to-human force of infection follows the usual mass action law and is the product of the number of mosquitoes per person, the daily mosquito biting rate and specific ZIKV infection transmissibility per day, the mosquito-to-human transmission rate [61], and the number *I^V^* of infected mosquitoes. Exposed mosquitoes transition to the infectious class at a rate *ε_V_* that is inversely proportional to the mean extrinsic latent period in the mosquito population [2]. Susceptible, exposed, and infectious mosquitoes all die at a rate that is inversely proportional to the mosquito lifespan, *μ_V_* [62]. The mosquito-to-human force of infection follows the usual mass action law. A full description of the stochastic model and the equations is provided in the Supplementary Materials.

**Fig. 7:**
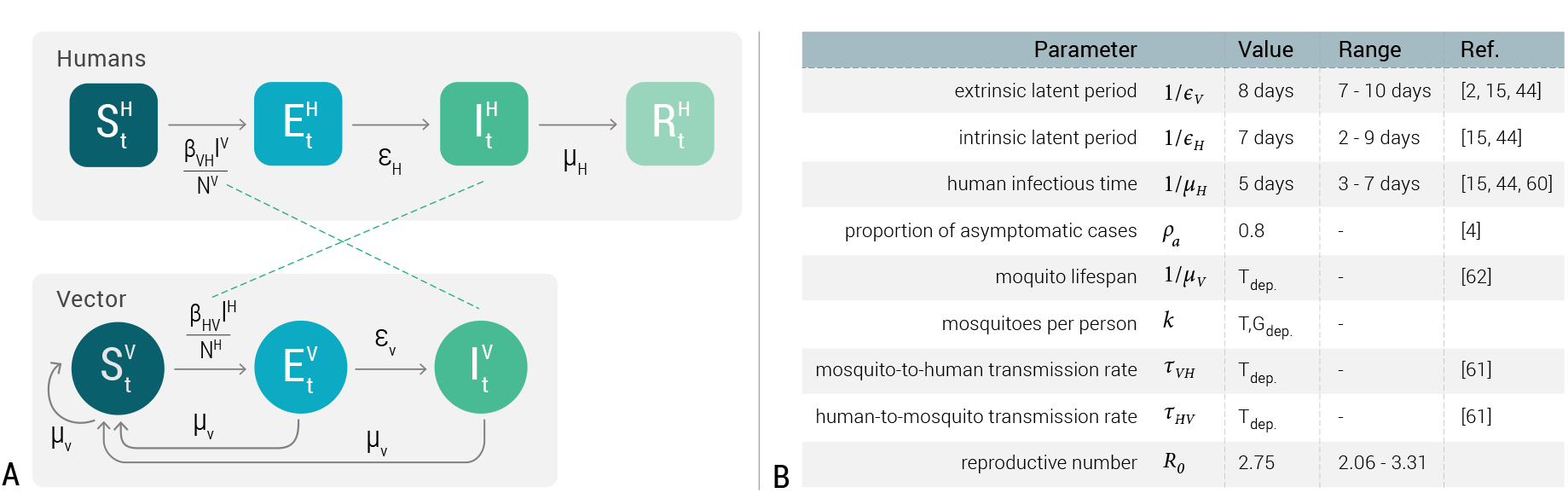
A) Compartmental classification for ZIKV infection. Humans can occupy one of the four top compartments: susceptible, which can acquire the infection through contacts (bites) with infectious mosquitoes; exposed, where individuals are infected but are not able yet to transmit the virus; infectious, where individuals are infected and can transmit the disease to susceptible mosquitoes; and recovered or removed, where individuals are no longer infectious. The compartmental model for the mosquito vector is shown below. B) Summary of the parameters of the model. *T_dep_*. denotes parameters that are temperature dependent. *T*, *G_dep_*. denotes parameters that are temperature and geo-location dependent. Specific values for the parameters can be found in Refs. [2, 60, 15, 4, 62, 61, 44].

A summary of the parameters defining the disease dynamic is reported in Fig. 7B. The empirical evidence related to the ZIKV disease in both human and mosquito populations is fairly limited at the moment. We have performed a review of the current studies of ZIKV and collected plausible ranges for these parameters. As in other studies, we have assumed that the drivers of ZIKV transmission are analogous to those of DENV. In particular, we have considered that mosquito lifespan, mosquito abundance, and the transmission probability per bite depend on the temperature level.

### Model calibration

The calibration of the disease dynamic model is performed by a Markov chain Monte Carlo (MCMC) analysis of data reported from the 2013 ZIKV epidemic in French Polynesia [15]. Setting the extrinsic and intrinsic latent periods and the human infectious period to reference values and using the average temperature of French Polynesia, we estimate a basic reproduction number (*R*_0_) of 2.75 (95% credible interval [2.53-2.98]), which is consistent with other ZIKV outbreak analyses [15, 63]. As *R*_0_ depends on the disease serial interval, we report a sensitivity analysis in the Supplementary Materials considering the upper and lower extremes of plausible serial intervals. Briefly, the estimated *R*_0_ values are 2.06 (95% credible interval [1.91-2.22]) and 3.31 (95% credible interval [3.03-3.6]) for the shortest and longest serial intervals, respectively. The *R*_0_ values are consistent with those estimated from local outbreaks in San Andres Island (*R*_0_ = 1.41) and Girardot, Colombia (*R*_0_ = 4.61) [64]. The calibration in French Polynesia provides the basic transmissibility of ZIKV. However, variations in temperature, mosquito abundance and other factors yield varying *R*_0_ in each subpopulation tracked by the model as discussed in the Supplementary Materials.

## Acknowledgements

We acknowledge funding from the MIDAS-National Institute of General Medical Sciences U54GM111274, the European Commission Horizon2020 CIMPLEX 641191 and to the Colombian Department of Science and Technology (Fulbright-Colciencias scholarship to D.P.R.)

